# A novel bioreactor technology for modelling fibrosis in human and rodent precision-cut liver slices

**DOI:** 10.1101/331173

**Authors:** Hannah L Paish, Lee H Reed, Helen Brown, Mark C Bryan, Olivier Govaere, Jack Leslie, Ben S Barksby, Jeremy French, Steven A White, Derek M Manas, Stuart M Robinson, Gabriele Spoletini, Clive Griffiths, Derek A Mann, Lee A Borthwick, Michael J Drinnan, Jelena Mann, Fiona Oakley

## Abstract

What is already known about this subject?

- Currently there are no effective anti-fibrotic drugs to treat liver fibrosis and there is an urgent unmet need to increase our knowledge of the disease process and develop better tools for anti-fibrotic drug discovery.
- Preclinical *in vitro* cell cultures and animal models are widely used to study liver fibrosis and test anti-fibrotic drugs, but have shortfalls; cell culture models lack the relevant complex cell-cell interactions of the liver and animal models only reproduce some features of human disease.
- Precision Cut Liver Slices (PCLS) are structurally representative of the liver and can be used to model liver fibrosis and test anti-fibrotic drugs. However, PCLS are typically cultured in elevated, non-physiological oxygen levels and only have a healthy lifespan of 48h.

What are the new findings?

- We have developed a novel bioreactor culture system that increases the longevity of functional PCLS to up to 6 days under normoxic conditions.
- Bioreactor cultured PCLS can be used to model fibrogenesis in both normal and fibrotic PCLS using a combination of biochemical and histological outputs.
- Administration of an Alk5 inhibitor effectively limits fibrogenesis in normal rodent and human PCLS and in rodent PCLS with established fibrosis.

How might it impact on clinical practice in the foreseeable future?

- The extended longevity of bioreactor cultured PCLS represent a novel pre-clinical tool to investigate the cellular and molecular mechanisms of liver fibrosis.
- Bioreactor cultured human PCLS offer a clinically relevant system to test efficacy of anti-fibrotic drugs.

**Abstract:** *Objective:* Precision cut liver slices (PCLS) retain the structure and cellular composition of the native liver and represent an improved system to study liver fibrosis compared to two-dimensional mono or co-cultures. The objective of this study was to develop a bioreactor system to increase the healthy lifespan of PCLS and model fibrogenesis.

*Design:* PCLS were generated from normal rat or human liver, or 4-week carbon tetrachloride-fibrotic rat liver and cultured in our patented bioreactor. PCLS function was quantified by albumin ELISA. Fibrosis was induced in PCLS by TGFβ1 and PDGFββ stimulation. Alk5 inhibitor therapy was used. Fibrosis was assessed by fibrogenic gene expression, Picrosirius Red and αSmooth Muscle Actin staining, hydroxyproline assay and collagen 1a1, fibronectin and hyaluronic acid ELISA.

*Results:* Bioreactor cultured PCLS are viable, maintaining tissue structure and stable albumin secretion for up to 6 days under normoxic culture conditions. Conversely, standard static transwell cultured PCLS rapidly deteriorate and albumin secretion is significantly impaired by 48 hours. TGFβ1 and PDGFββ stimulation of rat or human PCLS induced fibrogenic gene expression, release of extracellular matrix proteins, activation of hepatic myofibroblasts and histological fibrosis. Fibrogenesis slowly progresses over 6-days in cultured fibrotic rat PCLS without exogenous challenge. Alk5 inhibitor limited fibrogenesis in both TGFβ1 and PDGFββ stimulated PCLS and fibrotic PCLS.

*Conclusion:* We describe a new bioreactor technology which maintains functional PCLS cultures for 6 days. Bioreactor cultured PCLS can be successfully used to model fibrogenesis and demonstrate efficacy of an anti-fibrotic therapy.

## Introduction

Hepatic fibrosis is characterised by the accumulation of scar matrix in the liver and is the pathological consequence of persistent liver injury. Epithelial damage initiates local inflammation and activation of hepatic myofibroblasts (HM), which secrete extracellular matrix (ECM) proteins to form a temporary scar. If the injury stimuli desists, the scar is remodelled, however, persistent damage causes fibrosis[1].

Currently, liver cell mono or co-cultures, sandwich cultures, organoids or animal models are used to interrogate the mechanisms driving pathogenesis and reversion of liver disease and fibrosis to identify new targetable pathways and test anti-fibrotic drugs[2-4]. These preclinical tools have strengths but also limitations. Two-dimensional cell cultures lack the physiologically relevant cell:cell and cell:ECM interactions found in the intact liver and are exposed to supra-physiological levels of mechanical stress when cultured directly on plastic. The latter drastically alters cell behaviour; for example quiescent hepatic stellate cells (qHSC) grown on rigid tissue culture plastic trans-differentiate into αSMA+ HM that express pro-fibrotic genes and secrete high levels of ECM, whereas, culturing qHSC in soft matrigel retains the features of HSC quiescence[5]. Conversely, culturing HM on soft matrix hydrogels (∼2kPa) supresses pro-fibrotic gene expression and promotes a qHSC morphology[5, 6]. Hepatocytes rapidly dedifferentiate and down-regulate synthesis of albumin, metabolic enzymes and cytochrome P450 after 24 hours in culture[7]. 3D spheroids provide an alternative system to grow hepatocytes or mixed liver cell cultures but they fail to recapitulate the structural organisation of the liver or maintain physiologically relevant interactions with the ECM[4].

Numerous animal models have been described employing different injury stimuli to induce liver diseases[3], and study the events that drive fibrosis.. These models are mainly performed in young rodents, the disease inducer is often non-physiological, onset is rapid, there are metabolic differences between rodent and human liver and animals develop some but not all clinical features of the disease [3, 8].

Precision-cut liver slices (PCLS) are used as an ex-vivo culture model to study hepatic drug metabolism and fibrosis and benefit from retaining the 3D structure, physiological ECM composition and complex cell:cell interactions of the liver [9-11]. However, PCLS cultured under static conditions in normoxia typically have a limited functional lifespan of ∼24-48 hours due to hypoxia. This results in death and disruption of tissue architecture and dramatically reduces secretion of the functional marker albumin. Strategies to minimise the detrimental effects of hypoxia in PCLS and limit accumulation of metabolites include; increasing oxygen concentration from 20% to 40-95%, using synthetic oxygen carriers e.g. perfluorodecalin, varying media supplements/composition or introducing media flow by rocking or shaking the PCLS, using rotating culture vessels/rollers or perfusion circuits[10, 12, 13]. Recently, an air-liquid interface culture system has been used to model inflammation and immunological processes in human PCLS over a 15-day culture period, although of note this methodology was associated with significant spontaneous fibrogenesis[14]. A significant advantage of PCLS over animal models is that PCLS can be generated from human liver therefore liver disease biology, efficacy of therapies and drug metabolism can be modelled in human tissue *ex-vivo*[9, 15, 16].

We describe a new bioreactor technology that maintains the integrity and functionality of rodent and human PCLS while minimising tissue stress to enable inducible *ex-vivo* modelling of induced fibrogenesis and pre-clinical studies examining the efficacy of anti-fibrotic interventions.

## Materials and methods

### Animals

10-14 week old male Sprague Dawley rats were used in this study. Carbon tetrachloride (CCl_4_) treated rats received a 1:3 mixture of CCl_4_ and olive oil (0.1 mL per 100 g body weight, *via* intraperitoneal [IP] injection) biweekly for 4 weeks to induce liver fibrosis. Rats received humane care and experiments were approved by the Newcastle Animal Welfare and Ethical Review Board and performed under a UK Home Office licence.

### Human liver tissue

Human liver tissue was obtained from the normal resection margin surrounding colorectal metastasis. The tissue was obtained with informed consent from adult patients undergoing surgical resection at the Freeman Hospital, Newcastle-upon-Tyne, UK. This study was approved by the Newcastle & North Tyneside Research Ethics Committee (REC reference 12/NE/0395). Viability of hepatocytes in culture is significantly reduced when isolated from livers >3h post hepatectomy[17]. To minimise the ischaemic time and help preserve hepatocyte viability, the maximum time for generating PCLS for experiments was 2h post hepatectomy.

### Precision-Cut Liver Slices

Liver tissue was cored using a 8mm Stiefel biopsy punch (SmithKline Beecham, UK). Cores were transferred to a metal mould and submerged in 3% low gelling temperature agarose (A9414 Sigma-Aldrich, UK) and placed on ice for 2-5 minutes. Agarose embedded liver cores were super-glued to the vibratome mounting stage, submersed in the media chamber containing 4°C Hank’s Balanced Salt Solution+ and cut using a Leica VT1200S vibrating blade microtome (Leica Biosystems, UK) at a speed 0.3mm/sec, amplitude 2mm and thickness (step size) of 250μm (Fig.1A). PCLS were transferred onto 8µm-pore Transwell-inserts and cultured under static conditions in standard 12-well plates (Greiner Bio-one, Austria) or in a modified tissue culture plate (BioR plate) and rocked on the bioreactor platform (patent PCT/GB2016/053310) at a flow rate of 18.136 µl/sec (Fig.1B-C and Suppl Fig1A). For comparison, PCLS were cultured in Kirstall Quasivivo (qv500) units in Suppl Fig1B according to manufacturers instructions. All PCLS were cultured in Williams Medium E (W4128, Sigma-Aldrich) supplemented with 1% penicillin/streptomycin and L-glutamine, 1x Insulin Transferrin-Selenium X and 2% fetal bovine serum (Gibco, UK), 100nM dexamethasome (Cerilliant, USA) at 37°C supplemented with 5% CO_2_ and media were changed daily.

### Modelling fibrosis and anti-fibrotic drug therapy

PCLS were rested in culture for 24h then stimulated ± 3ng/ml TGFβ1 and 50ng/ml PDGFββ (fib stim) for a further 72h. For therapy PCLS were treated with 10μM Alk5 inhibitor SB-525334 for 1h prior to fib stim. Supernatants were collected at 24 hourly intervals and PCLS were harvested for histological and biochemical assays.

### Histology/immunohistochemistry

5-µm-thick formalin-fixed, paraffin-embedded (FFPE) liver sections were processed for haematoxylin and eosin (H&E),, Picrosirius Red and αSMA staining[18]. Images were acquired at x100 or x200 magnification using a Nikon ECLIPSE Ni-U microscope (NIS-Elements Br, Nikon, UK). The percentage Picrosirius red stained area was measured in fifteen-x200 magnification fields in Picrosirius red stained slides using Nikon Elements Imaging Software (NIS-Elements Br, Nikon, UK). The mean area was then calculated per slide.

### Enzyme-linked immunosorbent assay

ELISA quantifications for rat COL1a1 (LS-F11152, LSBio, UK), human (E88-129) and rat (E110-125) albumin (Bethyl laboratories, UK), human fibronectin (DY1918-05, R&D systems, UK) and human hyaluronic acid (DY2089, R&D systems, UK) were performed as per manufacturer’s instructions.

### Colorimetric assay

Aspartate aminotransferase (AST) levels were measured by the Clinical pathology department, Royal Victoria infirmary, Newcastle-Upon-Tyne, UK.

### Hydroxyproline assay

Two PCLS were weighed and digested with 1ml of 6N HCl to measure the quantity of hydroxyproline as previously described[19].

### RNA isolation, cDNA synthesis and PCR

2-4 PCLS per condition were placed in QIAzol, disrupted in a Qiagen Tissue Lyser II and passed through a Qiashredder (Qiagen, UK). Chloroform was added, the sample vortexed and centrifuged at 12,000g for 15 minutes. The aqueous layer was collected and added to 70% ethanol. Total RNA was purified using the RNeasy Micro Kit (Qiagen, UK) following manufacturers instructions. 1µg RNA was treated with 1µl DNase (Promega) for 30 minutes at 37°C and first strand cDNA produced *via* incubation with random hexamer primer (p(dN)6) and 100 units MMLV reverse transcriptase as previously described[20]. Real-time polymerase chain reaction was performed with SYBR Green JumpStart Taq ready mix (Sigma) as per manufacturer’s instructions using the primers listed in supplemental table 1 and relative level of transcriptional difference (RLTD) calculated using the 2^ΔΔCt^ method.

### Statistical Analysis

Data are presented as mean ± standard error of the mean (SEM), where *, **, *** and **** denote P values of <0.05, <0.01, <0.001 and <0.0001 respectively. All P values were calculated using GraphPad prism using an ANOVA with Tukey Post-hoc test.

## Results

### The bioreactor extends the lifespan of functional precision cut liver slices

Currently the healthy lifespan of static PCLS under normoxic conditions is reported to be around 48 hours due to rapid tissue degradation and loss of function[21]. To determine the optimal *in vitro* conditions for prolonged healthy maintenance of PCLS, we assessed and subsequently identified a number of critical parameters. To perform the tests, liver tissue was cored, embedded into an agarose block and 8mm diameter, 250μm thickness PCLS were prepared (Fig 1A) and cultured under a number of conditions. As an indicator of hepatocellular health and function within PCLS we employed production of albumin as an output. Initial pilot experiments demonstrated a need for continuous flow within the PCLS cultures. Therefore, we developed a bioreactor system consisting of a rocker and a BioR plate (Fig 1B and Suppl Fig 1A), which houses 6 bioreactor units, each unit comprising two cell culture wells connected *via* a channel (Fig 1B, right panel). When the BioR plate is rocked on the bioreactor platform (Suppl Fig 1A), media exchanges though the channel creating a bidirectional flow that helps oxygenate the slice and prevent accumulation of toxic metabolites. Culturing PCLS on transwell supports to prevent direct contact with plastic (Fig 1C). The transwell pore size was crucially important for PCLS viability, with a pore size of 8μm preserving albumin secretion at a stable level over 4 days (Fig 1D). Conversely, PCLS cultured in static BioR plate (i.e. without rocking) or on 3μm pore transwells (static or rocked) led to a rapid decrease in albumin production (Fig 1E). Similarly, rocking standard 12-well plates (without connecting channels between wells) or increasing the transwell insert pore size of static cultured PCLS from 3μm to 8μm did not improve hepatocellular function (Fig. 1E). We next asked if bidirectional flow was important and compared PCLS cultured in rocked BioR plates exposed to bidirectional flow, with PCLS cultured under unidirectional flow, in a circuit comprised either one or two connected Quasivivo units (Suppl Fig 1B). As expected, albumin production remained stable over 4 days in the bioreactor PCLS but decreased by ∼90% between days 2 and 3 under unidirectional flow (Suppl Fig 1B).

**Figure 1.**
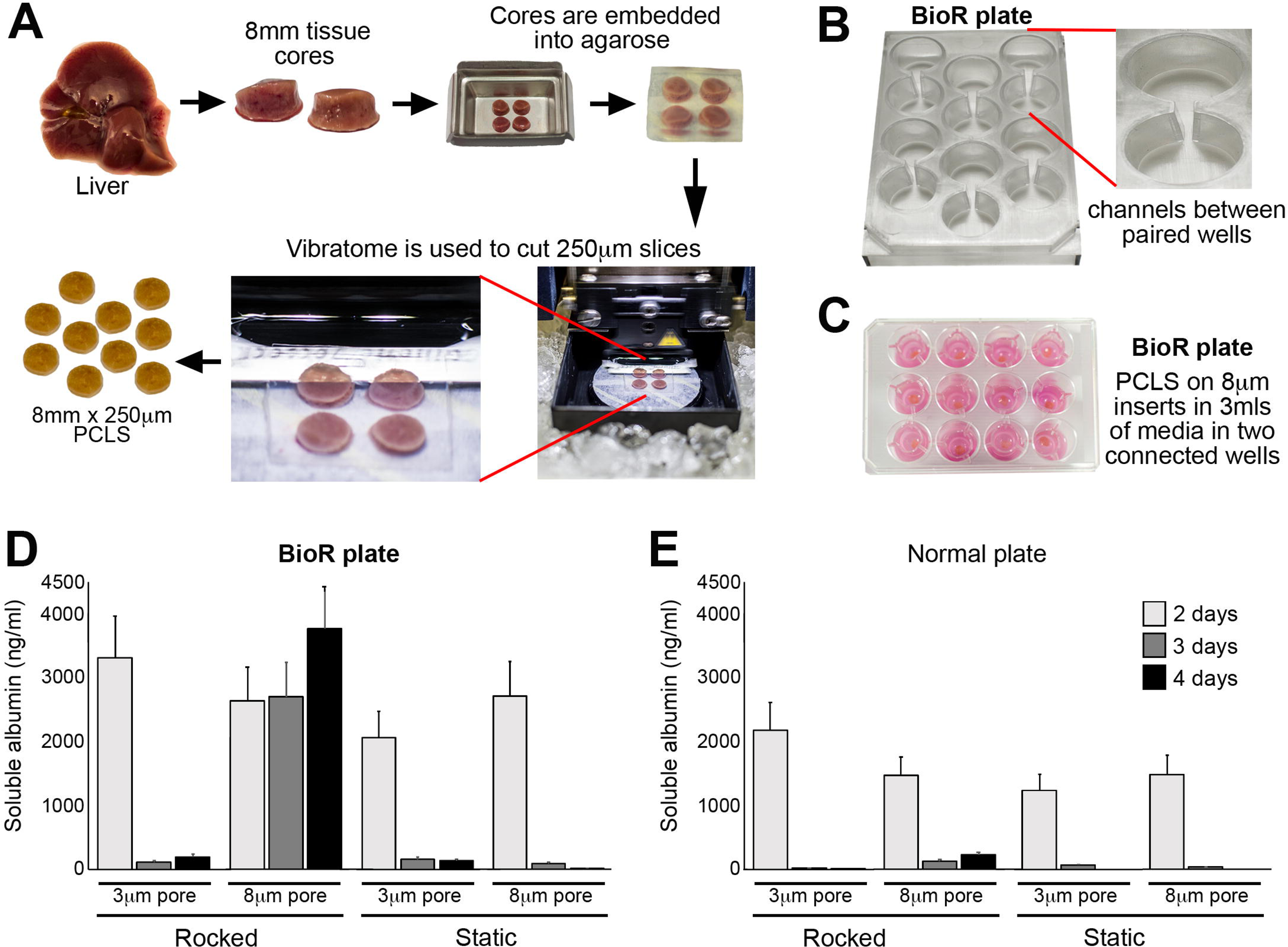
Generation of precision cut liver slices and optimization of the bioreactor culture method. (**A**) Workflow used to generate PCLS. (**B**) Photographs show the BioR plate containing 6 bioreactor chamber units and a zoomed image of one chamber unit, which contains two wells connected *via* a channel. (**C**) Photograph of PCLS cultured on 8μm transwell inserts in the BioR plate. (**D-E**) Graphs show secreted albumin (ng/ml) levels in the media of PCLS cultured on 3μm or 8μm transwell inserts in either a rocking BioR plate (**D**) or static 12-well transwell plate (**E**).

We next evaluated the histological appearance of PCLS cultured in the bioreactor or static transwell plates. Static cultured PCLS showed tissue disruption/degradation as early as 2 days, with loss of endothelial cells, atrophy of bile ducts, sinusoidal dilatation and loss of hepatocyte nuclei from days 2-6 days in culture (Suppl Fig. 2A). Conversely bioreactor-cultured PCLS retained structural integrity and the morphology of the portal tract and parenchyma (Suppl Fig. 2A). Consistent with the improved histological appearance of bioreactor cultured PCLS, liver albumin production was stable over the 6-day culture period. Conversely albumin levels were 33% lower in static cultured PCLS after 24h and significantly reduced over the next 5 days, reaching only 20-25% of the time-matched bioreactor-cultured PCLS (Suppl Fig. 2B). Collectively, these data confirm that culturing PCLS in our novel bioreactor under normoxic culture conditions and in the absence of oxygen carriers significantly increases the lifespan of functional liver slices compared to static transwell cultures.

**Figure 2.**
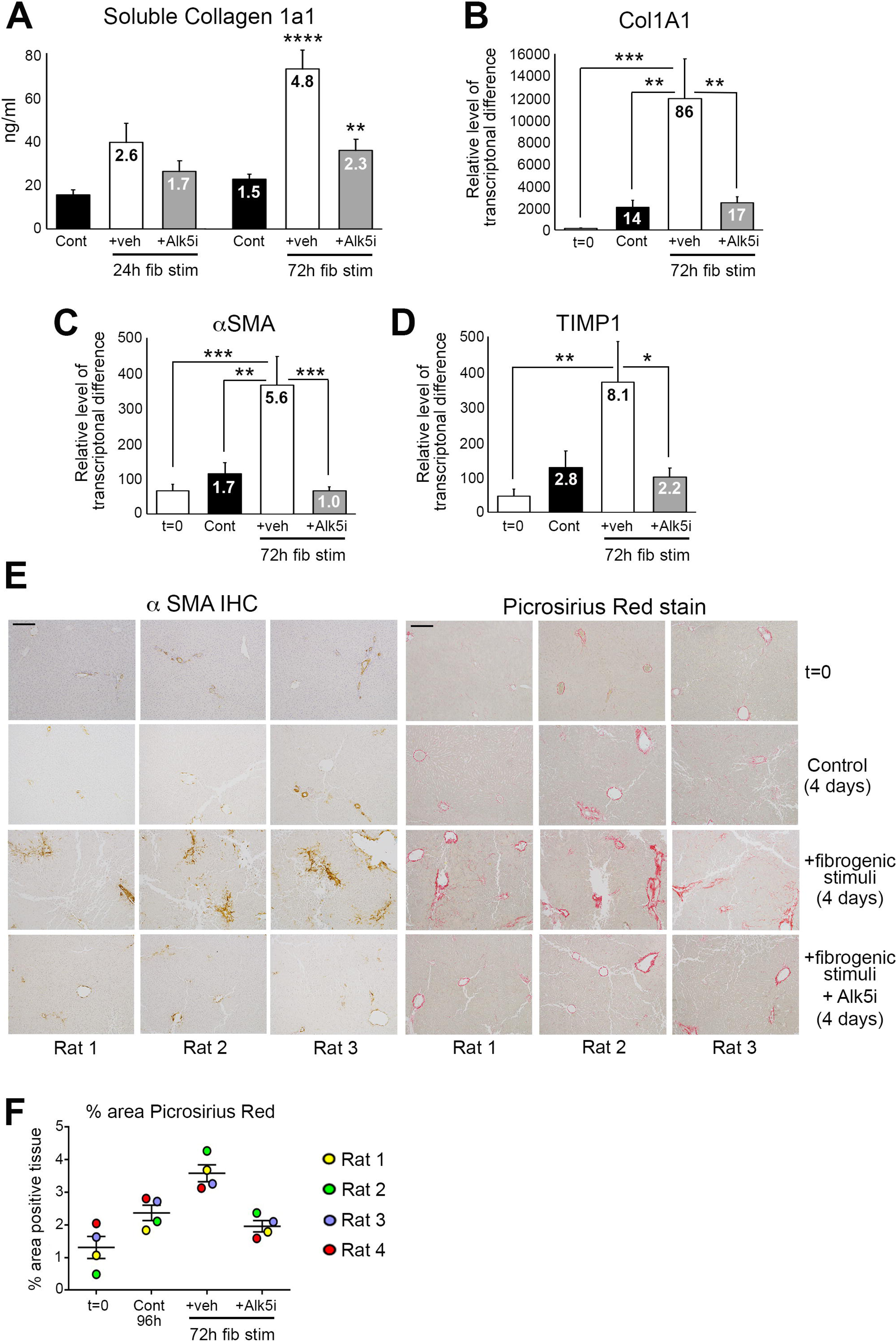
Modeling fibrosis and testing anti-fibrotic therapy in normal rat liver slices. (**A**) Graph shows average soluble collagen 1a1 (ng/ml) levels in the media of bioreactor-cultured rat PCLS after 24h rest and then 24 and 72h culture ± fib stim (TGFβ1/PDGFββ) ± Alk5i (48h and 96h total culture), numbers on bars show fold change compared to 48h control. (**B-D**) Graphs show mRNA levels of collagen 1A1, αSMA and TIMP1 in rat PCLS at t=0 and after 24h rest + 72h culture ± fib stim ± Alk5i (96h totalculture), numbers on bars show fold change compared to t=0. (**E**) Representative x100 magnification images of αSMA and Picrosirius red stained rat PCLS from three different livers at t=0 and after 24h rest plus 72h culture ± fib stim ± Alk5i. Scale bars equal 200 µm. (**F**) Graph shows the percentage area of picrosirius red stained tissue in rPCLS at t=0, and after 24h rest plus 72h culture ± fib stim ± Alk5i. Data are mean ± SEM in n=4 different livers. P values were calculated using Anova with Tukey post-hoc t-test (* P <0.05, ** P <0.01, *** P <0.001, *** P <0.0001).

The increased longevity of bioreactor-cultured PCLS that retain the cellular composition and tissue architecture provides the opportunity to induce fibrosis in the PCLS and assess efficacy of anti-fibrotic drugs. PCLS were generated from healthy rat liver, rested for 24h and then stimulated for up to 72h (up to 96h total culture period) with or without the profibrotic cytokine TGFβ1 in combination with platelet-derived growth factor-ββ (PDGFββ), which stimulates HM proliferation, migration and survival. After 96h culture basal soluble collagen secretion from control hPCLS into the culture media increases 1.5 fold, compared to 48h control slices (24h rest + 24h no treatment) (Fig. 2A). Conversely, soluble collagen 1a1 (COL1a1) secretion increased 2.6-fold after 24h and 4.8-fold after 72h TGFβ1/PDGFββ stimulation, which reached significance at 72h post-treatment (Fig. 2A). Col1a1 secretion was significantly attenuated by an Alk5 inhibitor (Alk5i), a potent supressor of TGFβ type I receptor dependant signalling[22]. Expression of pro-fibrotic genes col1a1, αSMA and tissue inhibitor of metalloproteinase 1 (TIMP1) were significantly elevated after 72h of TGFβ1/PDGFββ stimulation and this was supressed by Alk5i (Fig. 2B-D). Whilst αSMA and TIMP1 levels in 96h control PCLS were modestly increased compared to t=0 normal liver, collagen gene expression was significantly increased (14-fold) in control PCLS compared to t=0. However, important to note was that the increase in collagen gene expression did not translate to increased secretion of collagen 1a1 or to the deposition of fibrotic matrix within the PCLS tissue (Fig. 2A+E).

We next confirmed histologically that TGFβ1/PDGFββ promoted net fibrogenesis and deposition of fibrotic matrix in the PCLS (Fig. 2E). In t=0 healthy rat PCLS αSMA and Picrosirius red staining was restricted to vessels. Occasional αSMA+ HM were present in 4-day cultured control PCLS, whilst picrosirius red staining showed thickening of collagen around central veins and modest peri-sinusoidal collagen deposition (Fig. 2E). Conversely, TGFβ1/PDGFββ induced activation of αSMA+ HM and more extensive peri-sinusoidal fibrosis, with thick multi-layered collagen present around vessels. Alk5i blunted these histological changes such that αSMA and Picrosirius red staining was similar between control and TGFβ1/PDGFββ plus Alk5i PCLS (Fig. 2E). Collagen deposition, assessed by picrosirius red area analysis showed a small increase in collagen area in 4-day control PCLS compared to the donor matched t=0 PCLS. TGFβ1/PDGFββ stimulation significantly increased collagen deposition in all donors compared to t=0 and control cultured PCLS and this was significantly attenuated by Alk5i therapy (Fig. 4F). Importantly, AST and albumin release by the PCLS were not significantly different between the various experimental conditions, this suggesting that fibrotic stimuli or Alk5i therapy did not cause liver toxicity or effect hepatocellular function (Suppl Fig. 3A-B).

**Figure 3.**
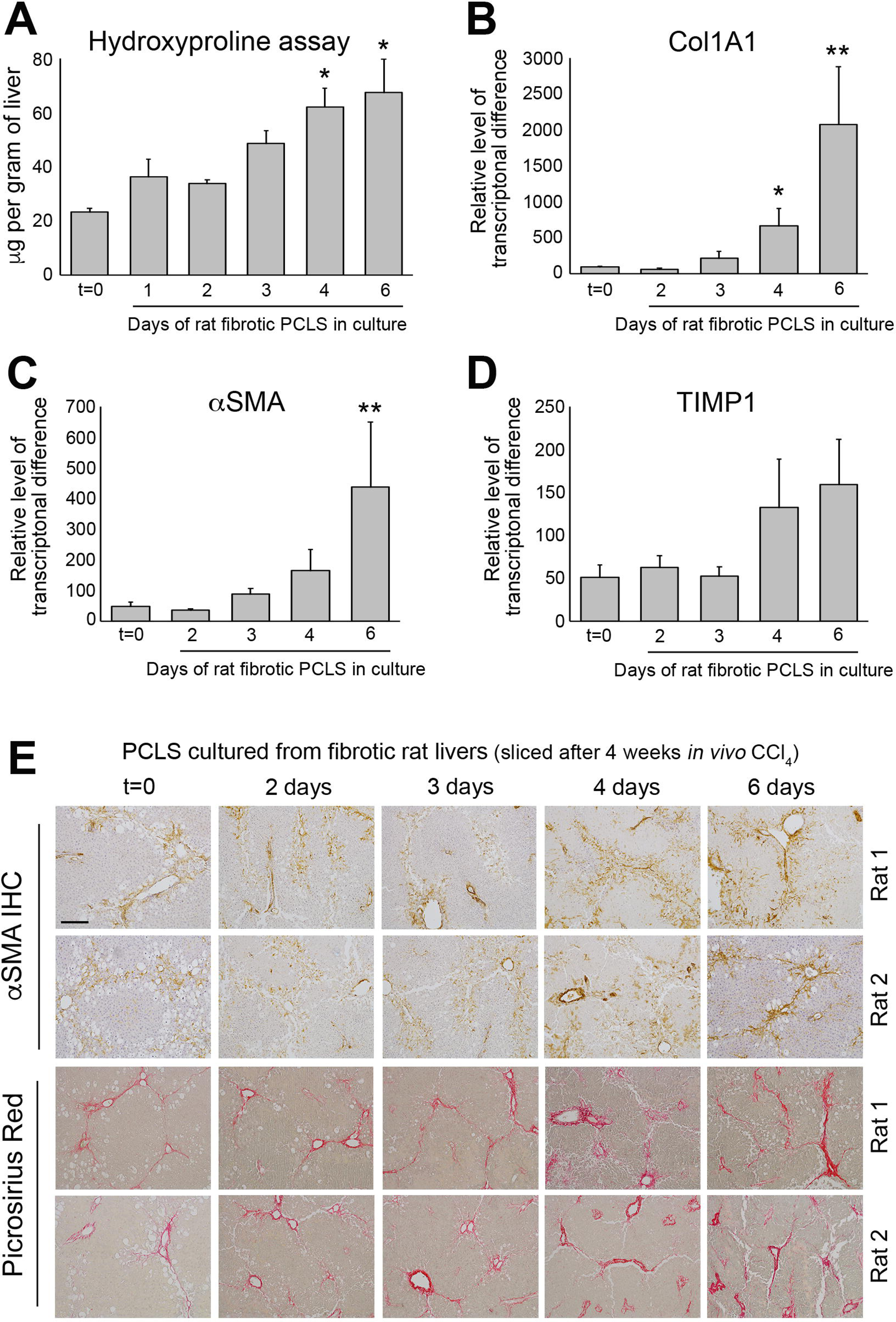
Bioreactor cultured fibrotic rat liver slices can be maintained for 6 days. (**A**) Graph shows hydroxyproline (μg/g of liver tissue) in bioreactor-cultured fPCLS. (**B**-**D**) Graphs show mRNA expression of collagen 1A1, αSMA and TIMP1 in bioreactor-cultured fPCLS at t=0 and 2, 3, 4- and 6-days culture. Data are mean ± SEM in n=5 independent slice experiments. (**E**) Representative images of αSMA and Picrosirius red stained rat fPCLS at t=0 and 2, 3, 4- and 6-days in culture. Scale bars equal 200µm. P values were calculated using an Anova with Tukey’s multiple comparisons test (* P <0.05 and ** P <0.01).

**Figure 4.**
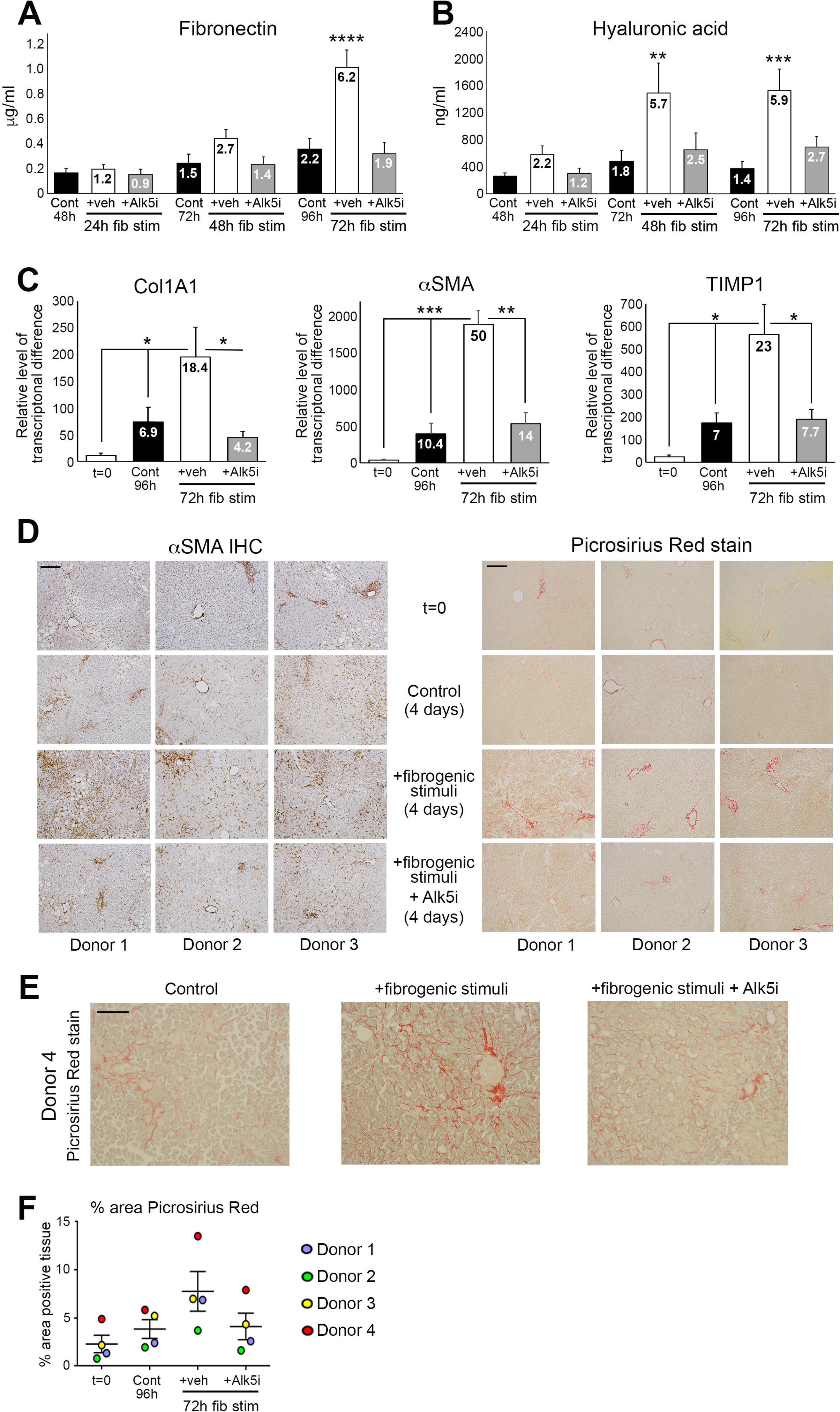
Modeling fibrosis and testing anti-fibrotic therapy in human liver slices. (**A-B**) Graphs show fibronectin (μg/ml) and hyaluronic acid (ng/ml) levels in the media of bioreactor-cultured hPCLS after 24 rest and then 24h or 72h culture ± fib stim (TGFβ1/PDGFββ) ± Alk5i, numbers on bars show fold change compared to 48h control. (**C**) Graphs show mRNA levels of collagen 1A1, αSMA and TIMP1 in hPCLS at t=0 and after 24h rest plus 72h culture ± fib stim ± Alk5i, numbers on bars show fold change compared to t=0. (**D**) Representative 100x magnification images of αSMA (left panel) and Picrosirius red (right panel) stained hPCLS from three different donor livers at t=0, and after 24h rest plus 72h culture (4-days) ± fib stim ± Alk5i. Scale bars equal 200 µm. (**E**) Representative 200x magnification image of Picrosirius red stained hPCLS form donor 4. (**F**) Graph shows the percentage area of picrosirius red stained tissue in hPCLS at t=0, and after 24h rest plus 72h culture ± fib stim ± Alk5i (96h total culture). Data are mean ± SEM in n=4 independent slice experiments. P values were calculated using an Anova with Tukey’s multiple comparisons test (* P <0.05, ** P <0.01, *** P <0.001 and **** P <0.0001).

We next explored the possibility to culture PCLS with pre-established fibrosis in the bioreactor and asked if the fibrosis is stable or dynamic. Fibrotic PCLS (fPCLS) were generated from 4-week carbon tetrachloride (CCl_4_) treated rats and cultured for up to 6 days. At t=0, H&E stained PCLS showed the expected features of CCl_4_ induced liver injury including peri-central damage and tissue necrosis, inflammation and ballooned hepatocytes (Suppl Fig. 4A). Consistent with a damaged liver, AST levels were high after 24h in culture, but decreased by 48h and continued to steadily decrease up to 6 days (Suppl Fig. 4B). After a 24h rest period, albumin production by fPCLS was slightly reduced but remained stable from days 2-6 (Suppl Fig. 4C). Levels of hydroxyproline, a major component of fibrotic matrix, were stable in fPCLS for 48h but then gradually increased with time, this effect reaching significance at 4 and 6 days, suggesting that ECM biosynthesis and deposition is progressive in cultured fPCLS (Fig. 3A). This effect was accompanied by a significant increase in Collagen 1a1 gene expression at day 4 and 6 (Fig. 3B). Similarly, mRNA levels of αSMA and TIMP1 increased at days 4 and 6 (Fig. 3C-D). Histologically, t=0 fPCLS revealed damaged hepatocytes, a characteristic “chicken wire” distribution of αSMA+ HM’s surrounding the central veins and bridging fibrosis (Fig. 3E). Consistent with the biochemical data, distribution of HM and bridging fibrosis is stable for 3 days however HM numbers and Picrosirius red stained area increases by 4-6 days in culture. Picrosirius red staining revealed a thickening of fibrous septa, increased intensity of collagen staining, presence of multilayered collagen and peri-portal fibrosis at days 4 and 6 (Fig. 3E). These data confirm that fPCLS are a dynamic system in which an active process of progressive net fibrogenesis occurs. We therefore next asked if fibrogenesis in fPCLS could be suppressed by Alk5i. Hydroxyproline concentration was increased 2-fold in 4-day cultured fPCLS compared to t=0 and this was significantly attenuated by Alk5i treatment (Suppl Fig. 4D).

An advantage of PCLS is the possibility to model fibrogenesis and assess anti-fibrotic therapies in human liver tissue. Therefore we asked if fibrotic changes can be induced in bioreactor-cultured human PCLS (hPCLS) and if an Alk5i could limit this. hPCLS were cultured for a 24h “rest period” and then treated ± TGFβ1/PDGFββ ± Alk5i for up to 72h (up to 96h total culture). In the absence of fibrotic stimuli, culturing PCLS for 96h induced a small increase in secretion of the soluble ECM components fibronectin (2.2-fold) and hyaluronic acid (HA) (1.4-fold) into the culture media compared to 48h control PCLS (Fig. 4A-B). Conversely, 72h TGFβ1/PDGFββ treatment significantly increased fibronectin (6.2-fold) and HA (5.9-fold) secretion compared to control PCLS (Fig. 4A-B). This was accompanied by an elevation in the gene expression of Col1A1, αSMA and TIMP1 (Fig. 4C). Compared to t=0, Col1A1, αSMA and TIMP1 expression was increased by 18.4, 50 and 23 fold respectively in TGFβ1/PDGFββ treated hPCLS. Alk5i suppressed hPCLS secretion of ECM components and fibrosis gene expression (Fig. 4A-C). It has been reported that in human liver αSMA+ cells are present in the portal tracts, the peri-venular space and liver lobule with discontinuous staining of qHSC in adult human liver biopsies or surgical resections[23]. Consistent with this report, αSMA+ cells were detected in the central veins, portal tracts and peri-sinusoidal spaces of normal donor liver (Suppl Fig. 5). Picrosirius red stained collagen was predominantly restricted to the vessels with delicate sinusoidal staining in non-sliced donor livers (Suppl Fig. 5). The histological appearance of αSMA and picrosirius red in t=0 post slice hPCLS was consistent with the non-sliced human liver suggesting that the preparation method and physical process of slicing did not promote acute HM activation or fibrotic changes (Fig. 4D). Treatment of hPCLS with fibrotic stimuli promoted the accumulation of αSMA+ HM throughout the parenchyma and adjacent to the central veins and portal tracts (Fig. 4D). Early fibrotic changes in hPCLS cultured with fibrotic stimuli were evident by the presence of more intense and extensive peri-sinusoidal picrosirius red staining as well as the thickening and multi-layering of collagen around the vessels. Fibrotic stimuli induced HM activation and collagen deposition was attenuated by an Alk5i and the histological appearance was similar to 96h control PCLS, showing a small increase in HM activation and modest persinusoidal collagen deposition compared to t=0 (Fig. 4D-E). Picrosirius red area analysis showed a small increase in collagen area in 4-day control hPCLS compared to the donor matched t=0 PCLS. TGFβ1/PDGFββ stimulation further increased collagen area in all donors and this was blunted and comparable to control levels after Alk5i therapy (Fig. 4F). Importantly, stimulation of hPCLS with TGFβ1/PDGFββ ± ALK5i did not significantly affect albumin production (Suppl Fig. 6).

## Discussion

In this report we describe a novel bioreactor culture system that extends the health and functional longevity of PCLS to at least 6-days and importantly in the absence of obvious hepatocellular stress and significant spontaneous fibrogenesis. Moreover we report that active induced fibrogenesis can be modelled using TGFβ1/PDGFββ stimulation and that an Alk5 inhibitor attenuates this process. Importantly, to evaluate fibrogenesis in our PCLS cultures we have quantified multiple biological outputs including secretion of ECM components, changes in fibrogenic gene expression and tissue collagen (hydroxyproline assay) as well as histological changes. Notably, the fibrogenic outputs can be reproducibly measured in PCLS stimulated with TGFβ1/PDGFββ within a relatively short time frame (4 days) thus providing a rapid model for testing potential anti-fibrotic drugs in high quality, functional human liver tissue. Furthermore, the bioreactor supports extended culture of fibrotic liver that becomes self-sustainable in culture, but remains modulatable using an Alk5 inhibitor. Critically the ability of novel drugs to block fibrogenesis in PCLS can be compared to effective inhibition mediated by Alk5i.

Numerous published studies describe different methodologies to culture human and/or rodent PCLS for modelling liver toxicity, investigating the effects of alcohol, studying fibrosis or evaluating anti-fibrotic drug efficacy[11, 24-26]. However, several critical differences between the PCLS culture conditions in other studies compared to our bioreactor are noteworthy. Previous studies have cultured PCLS directly on plastic, where their longevity and functionality either is severely limited (to <48h) or prolonged PCLS longevity to up to 72h by culturing in high oxygen concentrations with or without gentle shaking in a circular motion[24, 25]. Whilst most culture systems use fully submerged PCLS, a recent study described a rocked platform where hPCLS are cultured on organoid transwell plates in an air-liquid interface in normoxia for up to 15 days[14]. The air-liquid interface system cultures PCLS that are exposed to air on the upper side and are submerged in media on the lower side. This alternative system was used to evaluate the immunological responses to viral and bacterial products in PCLS at a gene expression level [14] but of note the authors reported substantial fibrogenesis which prevented the model from being used to study induced fibrosis. By contrast our bioreactor was specifically designed to minimise hepatocellular damage to limit background fibrogenesis and thereby enable investigations of induced fibrosis. Three key features help the bioreactor to maintain functionality and extend the lifespan of viable PCLS. Firstly, the bioreactor uses a combination of rocking and gravity to promote media exchange between the two adjacent wells of the bioreactor chamber creating a bidirectional flow that helps minimise hypoxia and reduce accumulation of toxic metabolites within the PCLS. Media flow in our bioreactor system has been closely matched to the sinusoidal flow within intact liver (flow rate of 18.136 µl/sec). Secondly, PCLS are cultured on semi-permeable transwell inserts (8μm pores) that facilitate media exchange from above and below the PCLS and prevent direct contact with stiff tissue culture plastic. Thirdly, PCLS are cultured under physiologically relevant normoxic conditions. Our bioreactor uses gentle rocking to induce flow and therefore does not require a peristaltic pump. Whilst perfusion systems require multiple perfusion circuits to perform multiple treatments[27], our bioreactor system is quick and easy to assemble, with scope to scale up to many concurrently running bioreactors for large studies with multiple experimental parameters or compound dose-responses.

Our development of a hPCLS model of fibrosis is important as it provides a robust platform with which to strengthen pre-clinical confidence in potential therapeutic targets/compound for future clinical trials. For example compounds demonstrating anti-fibrotic effects in rodent fibrosis models and 2D/3D cell culture platforms could be screened in hPCLS for drug metabolism, toxicity and efficacy in human tissue. Moreover modelling fibrogenesis in human or rodent PCLS would reduce animal use in accordance with the 3Rs; reduction, replacement and refinement. Approximately 150-200 PCLS can be generated from a normal or diseased rat liver thus provides a unique opportunity to perform medium throughput drug screening/dose determination experiments in a reduced number of animals prior to *in vivo* studies. Compounds exhibiting therapeutic actions in 2D-cultures and rodent PCLS would be progressed to hPCLS for focused drug screening to gain important information about target engagement and efficacy in human liver tissue. These approaches could significantly reduce animal use and help to replace moderate and severe *in vivo* procedures. Another advantage of hPCLS is daily media sampling, which raises the possibility that clinically relevant soluble biomarkers can be measured longitudinally to monitor disease progression or assess drug efficacy. In our study we measured hyaluronic acid (HA) release from hPCLS as a marker of fibrogenesis and serum HA levels have been reported to correlate with fibrosis grades in patients with hepatitis C [28]. Whilst PCLS have significant advantages over cell-culture models, one limitation is the lack of infiltrating immune cells, which can modulate the disease process. A future development for our PCLS system will be to egress immune cells or subsets of immune cells into normal or diseased PCLS. This would provide an opportunity to investigate how different immune cell populations modulate disease biology and the fibrogenic process in intact human tissue.

Currently our bioreactor technology can be used to study TGFβ1/PDGFββ induced liver fibrosis and test anti-fibrotic drugs, however, in the future the utility of the system can be significantly increased by developing new disease models in hPCLS that faithfully recreate the features of different aetiologies of liver disease. Development of bespoke liver disease models in hPCLS could be employed to elucidate the disease-specific mechanisms promoting disease progression but also identify core mechanisms of disease pathogenesis and fibrosis. For example treating PCLS with disease relevant lipids and sugars could model non-alcoholic liver disease (NAFLD), whilst addition of lipids and bacterial products and/or immune cells could recreate features of advanced NAFLD and non-alcoholic steatohepatitis. Whereas, culturing PCLS with ethanol or alcohol metabolites, either with or without immune cells is a strategy to model alcoholic liver disease (ALD). The system also lends itself to assess drug metabolism and drug-induced liver toxicity or model liver failure using acetaminophen. Currently, 2D monocultures of hepatoceullar carcinoma (HCC) cell lines and xenograft or chemical induced animal models of liver cancer are primarily used to identify drugable biological process/pathways in HCC. Therefore addition of liver cancer cell-lines or patient-derived cancer cells to either normal or diseased-induced bioreactor cultured hPCLS could represent an improved system to study cancer growth or test anti-tumour therapies within the context of a 3D liver environment. This system could also be used to study immune cell interactions within the cancer microenvironment and test novel immunotherapies.

Establishment of improved pre-clinical models in our bioreactor-cultured PCLS to study liver disease and cancer would aid drug discovery and permit targeted testing of new therapeutics in clinically relevant disease models.

## Figure legends

**Supplementary figure 1- Bioreactor platform and culture plate.**

(**A**) Photographs show the bioreactor rocker platform and BioR plate. (**B**) Graph showing media albumin levels in PCLS cultured in either one or two connected Quasivivo units (unidirectional flow) or the BioR plate (bidirectional flow).

**Supplementary figure 2- Bioreactor cultured rat PCLS have an extended healthy lifespan compared to static transwell cultured PCLS.**

(**A**) Representative H&E images show 2, 4 and 6 day bioreactor and static cultured rat PCLS. Scale bars equal 200 µm. (**B**) Graph shows average levels of albumin (ng/ml) released from static and bioreactor cultured rat PCLS into the culture media. Data represents mean ± SEM in n=4 independent slice experiments.

**Supplementary figure 3- Viability and function of rat liver slices are not affected by profibrotic stimuli or an Alk5 inhibitor.**

(**A-B**) Graphs show albumin (ng/ml) and aspartate aminotransferase (IU/L) levels in the culture media of bioreactor cultured rat PCLS after 24 rest and then 24h-72h culture ± fib stim (3ng/ml TGFβ1 and 50ng/ml PDGFββ) ± Alk5i. All data are presented as mean ± SEM in n=4 independent slice experiments.

**Supplementary figure 4- Functional characterisation of fibrotic rat liver slices.**

(**A**) Representative 100x images of H&E stained sections from n=4 independent t=0 fibrotic rat PCLS. (**B-C**) Graphs show aspartate aminotransferase (IU/L) and albumin (ng/ml) levels in the culture media of fibrotic rat PCLS bioreactor cultured for 6 days. All data are presented as mean ± SEM in n=4 independent slice experiments. (**D**) Graph shows hydroxyproline in 4-day bioreactor-cultured fPCLS ± 10ng/ml Alk5i. Data are mean ± SEM in n=3 independent slice experiments. P values were calculated using an Anova with Tukey’s multiple comparisons test (** P <0.01).

**Supplementary figure 5- Picrosirius red and αSMA staining in normal human liver tissue**

(**A**) Representative 100x images of αSMA and Picrosirius red stained sections from unsliced normal human liver tissue from the normal margin of a colorectal cancer resection. Images represent six different donor livers.

**Supplementary figure 6- Functional and histological characterisation of human liver slice treated with profibrotic stimuli**

(**A**) Graph show albumin (ng/ml) levels in the culture media of bioreactor cultured human PCLS stimulated ±fib stim ± Alk5i. All data are presented as mean ± SEM in n=4 independent slice experiments.

